# A mitochondrial tipping point couples early hyperexcitability to late-stage failure in patient-derived ALS motor neurons

**DOI:** 10.64898/2026.03.20.713134

**Authors:** Jovan Prerad, Thomas Vanwelden, Marit van Gorsel, Seppe Vansteenkiste, Karolien Pipeleers, Marlies Reusen, Ilse Bastiaens, Thomas Libotton, Katrien Princen, Floor Stam, Gerard Griffioen, Marc Fivaz

## Abstract

Amyotrophic lateral sclerosis (ALS) is a motor neuron (MN) disease characterized by profound alterations in energy metabolism and progressive degeneration of MNs. Evidence from patients and model systems point to MN hyperexcitability as an early hallmark of ALS. How altered electrical activity intersects with energy metabolism, however, remains largely unexplored. To directly examine this relationship, we performed long-term longitudinal recordings of neuronal firing and mitochondrial function in patient-derived MNs harboring the pathogenic TDP-43 mutation A382T, together with their isogenic controls. A382T MNs displayed an early, transient phase of hyperexcitability peaking around 35 days in culture, which was also observed in MNs carrying a different TDP-43 disease variant (M337V). This hyperexcitable phase coincided with elevated mitochondrial function (hyperpolarized membrane potential and accelerated electron flow across respiratory complexes) and was ultra-sensitive to mild Fo/F1 ATPase inhibition, revealing near maximal mitochondrial output to meet increased energy demands. This phase was followed by a sharp decline in A382T MN firing frequency, mitochondrial depolarization, and the loss of approximately 50% of firing-competent neurons. Acute bidirectional manipulations of MN firing rates elicited homeostatic mitochondrial responses, further demonstrating tight coupling between neuronal activity and mitochondrial metabolism. Together, these findings uncover a pathological trajectory in which early hyperexcitability drives mitochondrial hypermetabolism, with progressive erosion of mitochondrial capacity, ultimately leading to late-stage MN failure.

## INTRODUCTION

Brain function and energy metabolism are deeply intertwined. Neurons consume substantially more energy than other cell types to restore ion gradients after synaptic and action potentials^1^ and to sustain neurotransmission^2^. Because nerve cells do not have significant reserves of high-energy molecules, they must synthesize ATP on demand by glycolysis or oxidative phosphorylation. The brain is therefore a metabolically vulnerable organ uniquely sensitive to fuel availability, and neurons are particularly susceptible to even small ATP deficits, with potential consequences for neurodegeneration^3^. To meet energy demands imposed by neuronal activity, neurons dynamically adjust their metabolic rate, through an array of signaling pathways that often converge on regulation of the mitochondrial electron transport chain (ETC) and the Fo/F1 ATP synthase^4–8^.

Coupling between neuronal activity and mitochondrial energy metabolism has thus far mostly been studied under non-pathological conditions^4,5,7–9^. Less is known about this relationship in neurodegenerative conditions, although aberrant neuronal activity and mitochondrial dysfunction are typical hallmarks of these diseases. Notably, the extent to which the interaction between neuronal activity and mitochondrial metabolism contributes to disease progression has rarely been addressed.

Amyotrophic lateral sclerosis (ALS) is a prime example of a neurodegenerative disease with profound alterations in neuronal firing and energy metabolism. It is a fatal condition characterized by the selective and progressive loss of cortical and spinal motor neurons (MNs), leading to paralysis and death within a few years of symptom onset^10–12^. Cortical hyperexcitability is now considered a central feature of ALS and is observed in both sporadic and familial ALS patients^13^, typically before disease onset^14^. MN hyperexcitability has also been reported in ALS mouse models harboring disease-causing mutations in prominent familial ALS genes, including *SOD1*, *FUS*, *C9orf72,* and *TARDBP*^15–19^. Riluzole, one of the few approved treatments for ALS, dampens neuronal activity by reducing glutamatergic neurotransmission, further highlighting the clinical relevance of early MN hyperexcitability^20,21^.

Modeling familial forms of ALS in patient iPSC-derived spinal MNs, the principal induced MN subtype generated in culture, also showed evidence for hyperexcitability, particularly in young neurons^22–25^, with conflicting results for SOD1 (G93A) MNs^26^. Culturing MNs with *TARDBP* or *C9orf72* ALS variants for longer time periods appears to render them hypoexcitable^23,24^, suggesting a pathological transition towards neuronal decline. One limitation of these iPSC-MNs studies is that changes in activity patterns were deduced from isolated electrophysiological measurements with limited longitudinal resolution and were not recorded in the same individual neurons over time. The mechanisms underlying MN hyperexcitability in these experimental systems are multifaceted and depend on the ALS gene being modeled, but they often lead to a shared outcome: elevated intrinsic MN excitability, reflected by a lower current threshold for action potential firing and increased spontaneous firing frequency^13^.

In parallel to these changes in MN activity patterns, ALS is associated with severe alterations in energy metabolism at the organismal, brain, and cellular levels^27^. Energy expenditure consistently exceeds intake in both sporadic and familial ALS patients^28^ and hypermetabolism (leading to weight loss) is associated with worse prognosis^29^. Accordingly, imaging studies in ALS subjects have identified brain regions with increased metabolic activity^30^. These system-level perturbations in metabolism are associated with profound changes in cellular metabolic pathways observed across disease model systems^31–35^, with evidence for widespread mitochondrial defects^36–41^.

Mitochondrial dysfunction is observed in nearly all transgenic and overexpression models of familial ALS and include defects in the electron transport chain (ETC)^36,39^, suboptimal (uncoupled) respiration^42,43^, impaired calcium buffering^37^, elevated production of reactive oxygen species (ROS),^42^ and exaggerated opening of the permeability transition pore^37,40^. It is worth noting, however, that mitochondrial defects appear far less penetrant in human iPSC– or fibroblast-derived ALS models^44–47^, presumably because of lower (endogenous) expression of heterozygous disease variants. Of interest, mitochondrial hypermetabolism has also been described in fibroblasts from sporadic and familial ALS patients^47,48^ and in a knock-in valosin-containing protein (VCP) model of ALS^40^, suggesting that the pathological signature of MN firing may be mirrored by corresponding changes in mitochondrial metabolism and function.

To probe the precise temporal relationship between MN firing and mitochondrial activity, we combined single-cell imaging, high-density multi-electrode arrays and ETC electron flow measurements to map the trajectory of neuronal and mitochondrial activity in human iPSC-MNs over 50 days in culture. We selected patient iPSCs carrying ALS mutations in *TARDBP,* since TDP-43 inclusions are found in 95% of sporadic ALS cases^49^, and TDP-43 ALS mutations are associated with abnormal MN firing patterns^19,23^ and a broad range of mitochondrial phenotypes^36,50,51^. We report co-evolution and interdependence of neuronal and mitochondrial activity in ALS MNs in a pathological trajectory that features early hyperexcitability/hypermetabolism and late-stage decline in neuronal and mitochondrial outputs. We developed a model that accounts for this biphasic neurometabolic signature based on accumulating damage of mitochondria functioning near their bioenergetic limit to maintain energy homeostasis. This model captures the course of ALS progression in patients –decades symptom-free followed by death 3-5 years after disease onset – and identifies a new area for therapeutics development at the intersection of electrical activity and energy metabolism.

## RESULTS

### Longitudinal profiling of neuronal firing reveals a shift from hyper– to hypo-excitability in TDP-43 (A382T) MNs

Patient-derived iPSCs harboring the heterozygous A382T TDP-43 mutation, one of the most common TDP-43 disease variants^52^, were differentiated into spinal MNs^53^, alongside their isogenic controls (A382A TDP-43) **(Supplementary Figure 1a)**. More than 80% of these cells express early markers of MN progenitors (ISL1 and HB9) and progressively acquire the characteristics of spinal MNs (ChAT+, NKX2.2– and GFAP-) **(Supplementary Figure 1b,c)**.

To monitor neuronal activity in individual MNs non-invasively, these cells were virally transduced with a genetically encoded calcium indicator (Neuroburst orange^TM^), at Day-in-Culture (DiC) 20, following dissociation of embryoid bodies, 30 days after neuralization of iPSCs by dual SMAD inhibition. Calcium spikes were imaged in MN cell bodies (3 Hz, 1 min epoch) at least once a day and as long as cultures remained viable, until DiC 35-66. In these conditions, calcium transients reflect bursts of spontaneous action potential firing in the MN network. Several firing parameters were extracted and quantified over time, including the number of active objects (neurons that fire at least once during the epoch), burst rate, burst duration, burst strength, and network correlation (synchronized activity). These metrics were measured in parallel in hundreds to thousands of A382T and control (ISO) MNs distributed across replicate wells of a 96-well plate (to minimize positional effects) and averaged across 5 independent cultures reaching DiC 50 to capture robust changes in firing patterns **(Figure 1)**. Alternatively, each independent MN culture was profiled separately, enabling analysis of firing behavior (for some of them) at later time points. **(Supplementary Figure 2a).**

**Figure 1.**
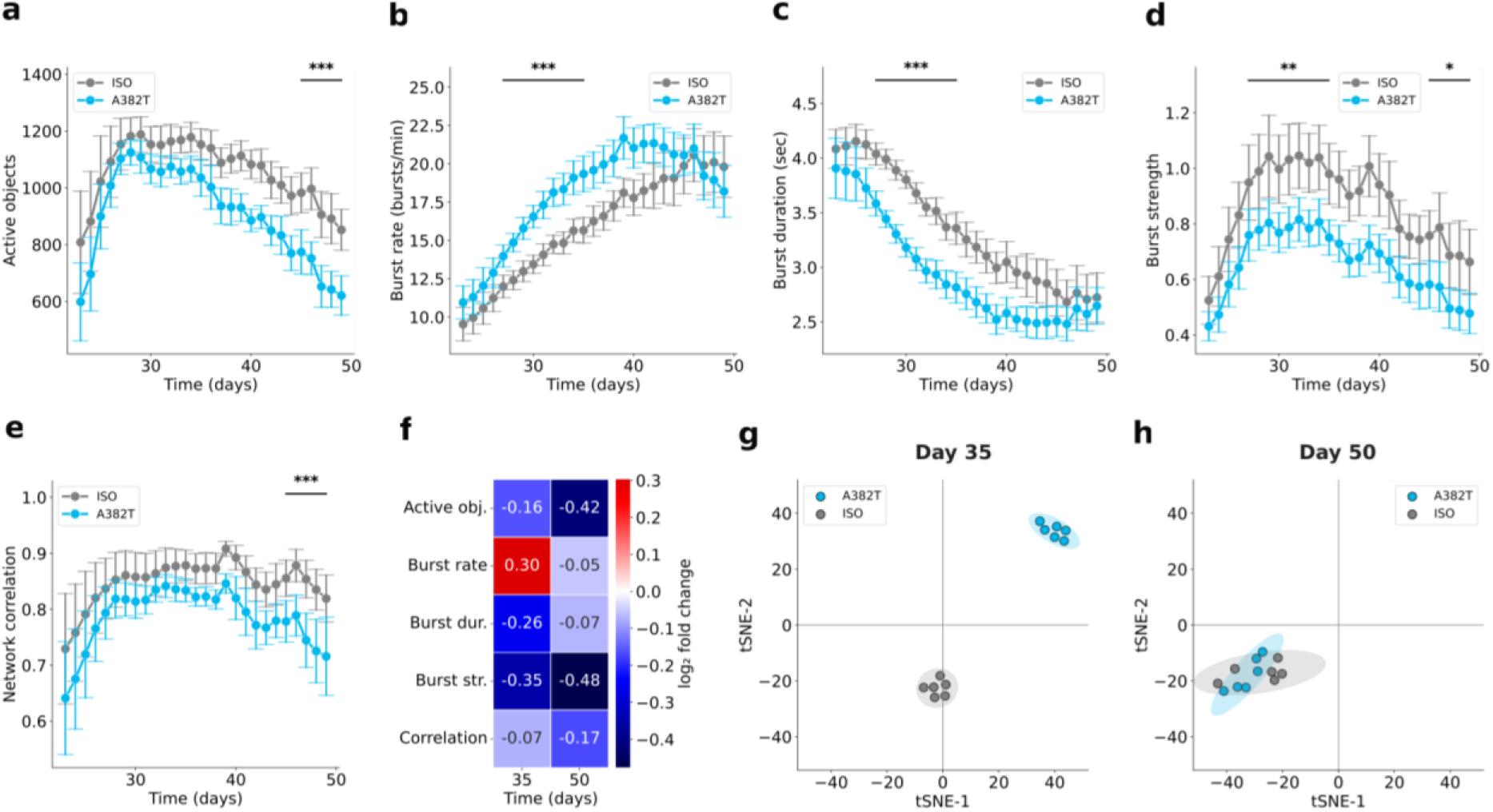
| Longitudinal profiling of Ca^2+^ transients reveals early hyperactivity and accelerated functional decline in A382T MNs. (**a–e**) Multi-dimensional analysis of spontaneous neuronal firing in A382T and ISO TDP-43 MNs from DiC 23 to 50. Time series showing firing metrics (mean ± SEM) from averaged wells originating from 7 independent MN cultures, with 5 cultures reaching the age of DiC 50. Statistics were performed using a linear mixed-effects model with genotype as fixed effect and experiment as random effect. *: p < 0.05, **: p < 0.01, ***: p < 0.001 indicate differences between ISO and A382T within early (DiC 27–35, n = 7) or late (DiC 45–50, n = 5) windows. **(f)** Heatmap showing mean A382T/ISO log2 fold change for each firing parameter at DiC 35 and 50. **(g–h)** t-SNE analysis of firing parameters from a representative experiment at DiC 35 and 50; dots indicate individual wells. Ellipses indicate the 95% confidence region of t-SNE embeddings for each genotype.

These MN cultures display high levels of spontaneous activity, regardless of genotype, a common feature of healthy iPSC-derived MN preparations. Up to DiC 35, similar active object counts and network correlations were measured in A382T and ISO MN cultures **(Figure 1a,e)** reflecting matching cell density, and network connectivity. From DiC 35 onwards, however, A382T MNs experienced a sharp decline in active object count compared to their ISO counterparts and a corresponding trend in network synchrony **(Figure 1a,e)**, indicating accelerated loss of firing-competent neurons. This likely reflects an impact on activity rather than cell survival, as we failed to detect clear signs of toxicity caused by expression of A382T TDP-43 **(Supplementary Figure 2c)**.

Analysis of firing frequency in active objects revealed an increasing and markedly higher burst rate in A382T MNs until DiC 40, at which point it stabilized and started to decline **(Figure 1b, Supplementary Figure 2a)**. ISO burst rates, in contrast, kept increasing with a “cross-over” between A382T and ISO trajectories at DiC 45 **(Figure 1b)**. The initial increase in burst rate observed in both groups is mirrored by a concomitant decrease in burst duration and likely reflects maturation of these cells^54^ **(Figure 1b,c)**. Burst strength was higher in the ISO MNs consistent with greater calcium entry during each burst, suggesting less calcium is needed for A382T MN firing **(Figure 1d)**. This longitudinal analysis of firing patterns revealed an initial phase of hyperexcitability in A382T MNs followed by progressive loss of active neurons and reduced network synchronization.

Comparison of these firing parameters at two time points: DiC 35 (hyperexcitable phase) and DiC 50 (declining phase) exemplified the switch in A382T MN firing patterns **(Figure 1f)**. We used t-distributed stochastic neighbor embedding (t-SNE) for global visualization of these firing properties. At DiC 35, firing parameters in A382T and ISO MNs separated into two distinct clusters **(Figure 1g)**, indicating a disease-specific firing signature of ALS MNs. These two clusters converged at DiC 50 **(Figure 1h, Supplementary Figure 2b)**, reflecting more similar firing patterns between the two groups in the declining phase.

To determine whether early hyperexcitability is observed in MNs harboring other TDP-43 ALS mutations, we recorded spontaneous neuronal activity in M337V TDP-43 MNs^52^, together with their own isogenic controls **(Supplementary Figure 1d)**. M337V MNs also exhibited higher burst rates relative to their isogenic counterparts, up to DiC 45 **(Supplementary Figure 2d,e)**. It was difficult to keep these two isogenic MN cultures alive beyond DiC 45, preventing us from studying their behavior at a later time point. Nonetheless, these findings suggest that early on, elevated MN firing frequency may be a general feature of MNs with familial TDP-43 mutations.

### Multi-electrode array recordings confirm early hyperexcitability and reveal intense burst propagation in A382T TDP-43 MNs

To examine firing patterns with increased temporal resolution, we grew A382T and ISO MNs on high density multi-electrode arrays (MEAs) and recorded electrical activity of the network at an early point (DiC 25) during the A382T hyperexcitable phase. Analysis of these recordings allowed us to distinguish single spikes (single electrode responses) from bursts and revealed a striking increase in firing activity in A382T MNs compared to their isogenic counterparts **(Figure 2)**.

**Figure 2.**
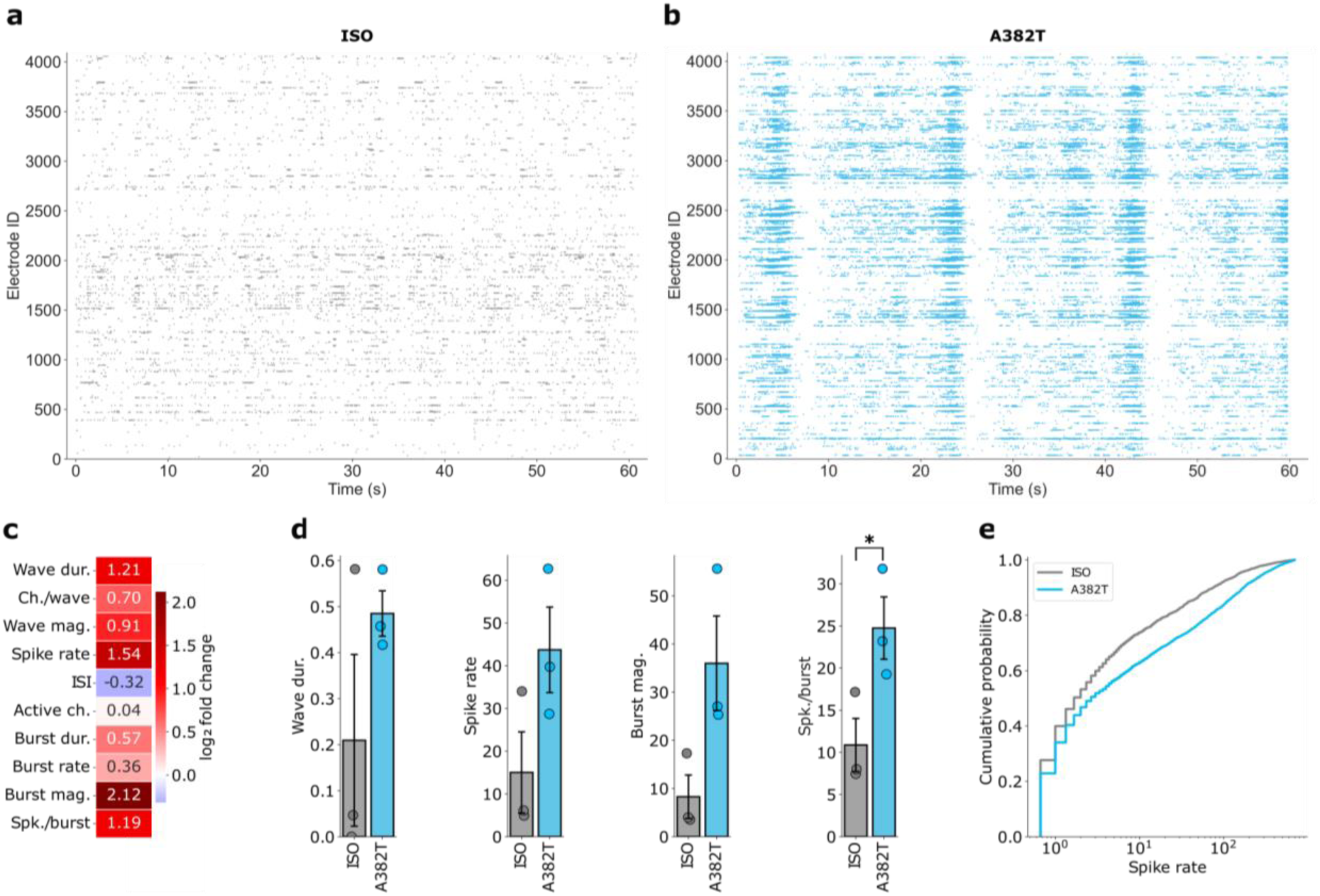
| MEA recordings reveal a global increase in firing behavior in A382T MNs at DiC 25. (**a,b**) MEA raster plots of ISO and A382T MNs at DiC 25, with each row representing the activity profile of a single electrode. **(c)** Heatmap representation of the average log_2_ A382T/ISO ratio for each MEA parameter, from three replicate MEA wells for each genotype. **(d)** Comparison of four firing parameters across genotypes shown as mean ± SEM. *: p < 0.05, Welsh’s t-test (n = 3 MEA wells). Data points represent the mean electrode value per MEA well. **(e)** Cumulative probability distribution of spike rates for ISO and A382T MNs, pooled across all electrodes (n = 4094) from three independent MEAs per genotype.

Hyperactivity in A382T MNs is immediately apparent in side-by-side comparison of raster plots, with the presence of burst waves that rhythmically propagate throughout the entire network **(Figure 2a,b)**. This network-wide oscillatory activity is surprisingly robust in TDP-43 mutant MNs **(Figure 2b)** and is less pronounced in ISO MNs, although some of these cells display coordinated rhythmic patterns **(Figure 2a).** The number of active channels was not significantly different between A382T and ISO MNs, reflecting similar electrode coverage and cell densities **(Figure 2c, Supplementary Figure 2e)**; all other firing metrics, however, point towards marked elevation of neuronal activity in A382T MNs **(Figure 2c)**, and include a higher spike rate with a concomitant diminution of the inter-spike interval **(Figure 2c,d)**. Burst metrics are all elevated in A382T MNs, with a >4-fold increase in burst amplitude, together with elevation of the number of spikes per burst, burst rate, and burst duration **(Figure 2c,d).** Likewise, burst wave magnitude, the number of channels per wave, and wave duration are all higher in A382T MNs. Although the average spike rate per MEA chip was not significantly different between A382T and ISO neurons, the cumulative distribution of spike rates across individual electrodes is significantly shifted toward higher spike frequencies in A382T MNs compared to ISO controls (Kolmogorov-Smirnov test, p = 4.930e-17) **(Figure 2e).** When grown on these MEA chips, MNs showed signs of toxicity after DiC 40, impeding MEA recordings at later time points. Collectively, these MEA recordings reveal firing behaviors largely concordant with those observed by calcium imaging **(Figure 1)** and confirm early and robust MN hyperexcitability induced by the pathogenic A382T TDP-43 mutation.

### Electron flow across respiratory complexes is transiently elevated in A382T MNs during the hyperexcitable phase

Electron flow across the ETC drives cellular respiration and ATP synthesis and is therefore a reliable indicator of mitochondrial metabolism. To measure ETC activity, we employed an electron-flow-based assay relying on ETC-driven reduction of a tetrazolium dye^55^ in semi-permeabilized MNs exposed to a customized array of energy substrates. These mitochondrial measurements were carried out in parallel to neuronal activity measurements in separate plates of the same MN cultures. At early time points (DiC 27-32), A382T MNs exhibited significantly higher electron flow compared to ISO MNs, in response to all tested substrates **(Figure 3a,c)**. The concurrent elevation of ETC activity in response to complex I (malate, pyruvate), and complex II (succinate) substrates reflects a global adaptive response rather than substrate– or complex-specific effects. At later stages (DiC 60), however, electron flow in A382T neurons converged to ISO levels, mirroring the decrease in neuronal activity **(Figure 3b,c)**.

**Figure 3.**
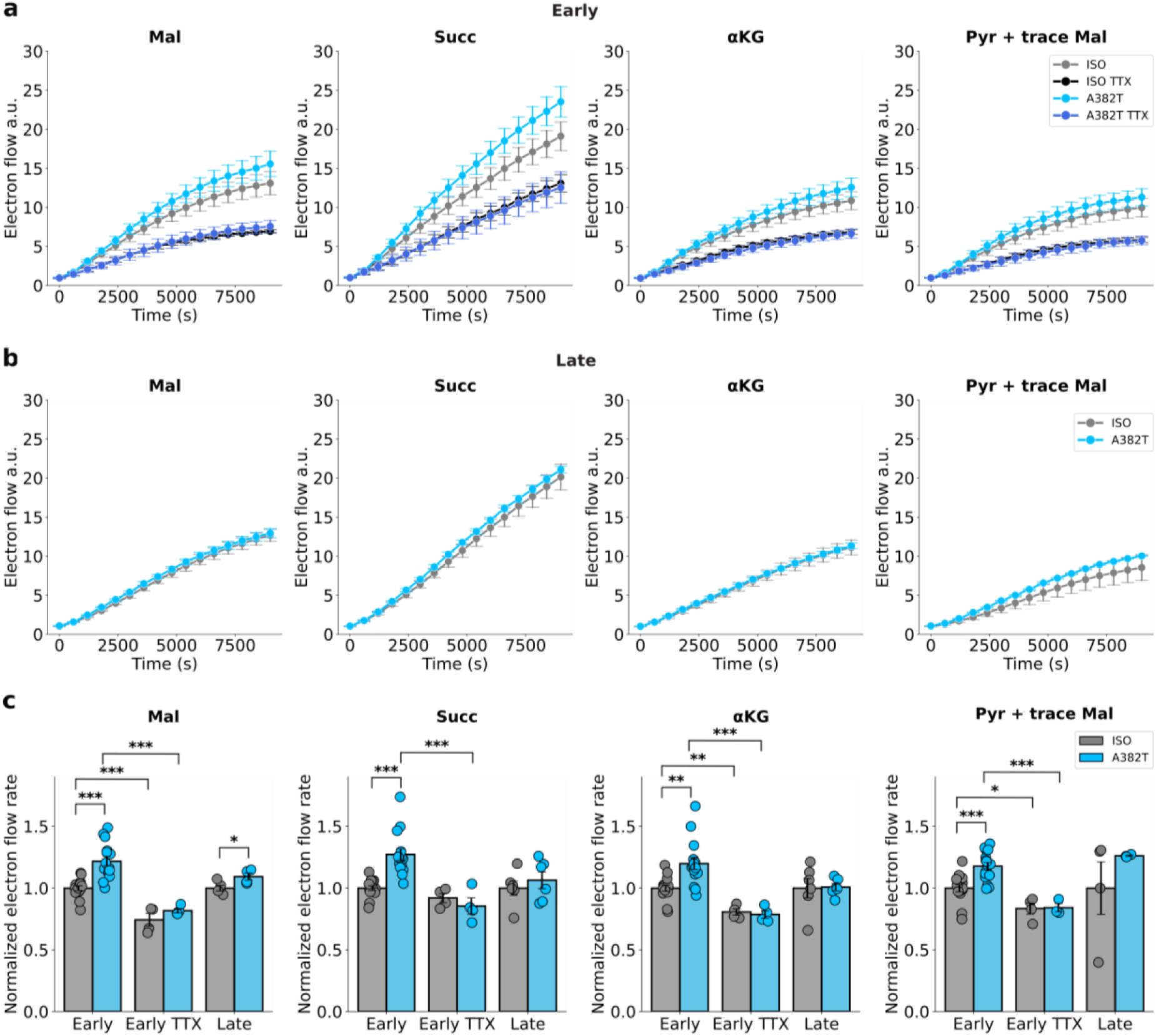
| A382T MNs show a transient elevation of ETC activity. (**a,b**) Kinetics of ETC electron flow fueled by the indicated substrates in A382T and ISO MNs at early (DiC 27-32, **a**) and late (DiC 60, **b**) time points, with or without TTX (1 µM, 24 h). **(c)** Quantification of electron flow rate in early-stage (DiC 27-32) MNs treated with TTX or vehicle and in older cultures (DiC 60). Data points represent individual wells derived from 5 (Early) or 2 (Early/TTX, Late) independent cultures. Bar plots show mean ± SEM. Statistical significance was determined using either an unpaired t-test or a Mann-Whitney U test for normally and non-normally distributed datasets (A382T Early vs Late Succ). *p < 0.05, **p < 0.01, ***p < 0.001.

Elevation of ETC activity in A382T MNs during the hyperexcitable phase suggests an adaptive mitochondrial response to sustain the increased ATP demand. To determine whether inhibition of neuronal activity is sufficient to reduce ETC activity, we blocked action potential firing with tetrodotoxin (TTX) in MN cultures at early time points (DiC 27-32) **(Supplementary Figure 3b)** and measured the impact of this perturbation on ETC activity 24 hours later. TTX induced a marked reduction in electron flow rates both in A382T and ISO MNs, demonstrating a clear influence of neuronal activity on mitochondrial metabolism **(Figure 3a,c).**

### Time-dependent reversal of mitochondrial membrane potential in A382T MNs with a tipping point at the peak of hyperexcitability

Next, we measured the mitochondrial membrane potential (ΔΨ_m_), a central component of the electrochemical gradient across the inner mitochondrial membrane (IMM), using the cationic fluorescent dye TMRM (tetramethyl rhodamine methyl ester)^56^. Analysis of TMRM fluorescence in individual organelles revealed elevated ΔΨ_m_ in A382T MNs at DiC 34 **(Figure 4a,b)**, consistent with increased ETC activity in the hyperexcitable state **(Figure 3a,c)**. At DiC 60, ΔΨ_m_ remained stable in ISO MNs, but dropped in A382T MNs **(Figure 4a,b)**, with no significant impact on mitochondrial count **(Supplementary Figure 3a)**, indicating depolarization and functional impairment of these organelles at end-stage.

**Figure 4.**
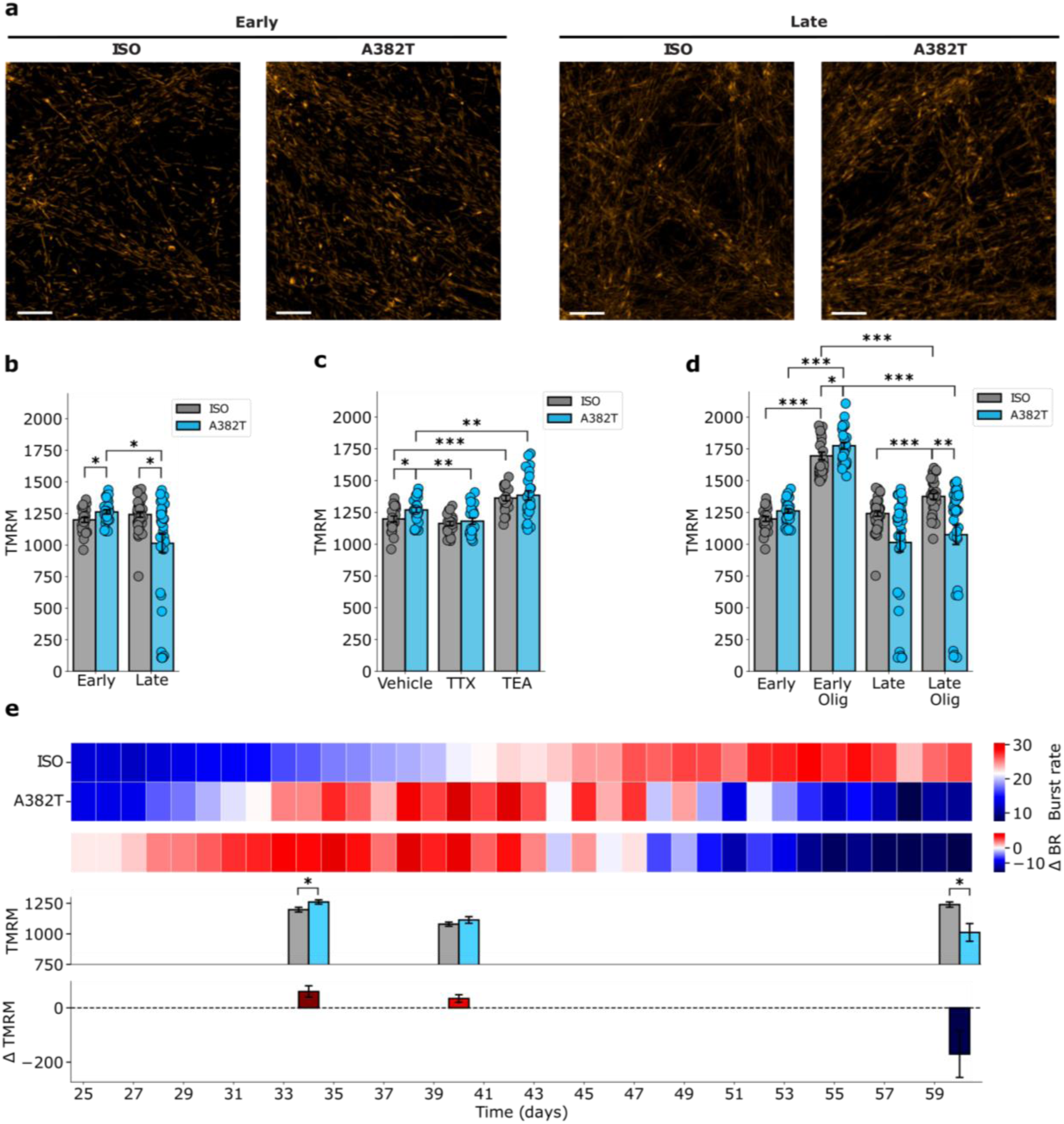
| Mitochondrial membrane potential in A382T MNs follows a biphasic trajectory. **(a)** DY_m_ imaged by TMRM staining in neurites of ISO and A382T MNs at early (DiC 34) and late (DiC 70) time points. **(b-d)** Mean TMRM intensity from individual mitochondria averaged across wells from four independent experiments in early (DiC 34) and late (DiC 60) neurites of ISO and A382T MNs. **(c)** Influence of activity modulators TTX (1 µM) and TEA (4 mM) on DY_m_ 24 h after treatment in early-stage MN cultures. **(d)** Impact of oligomycin (10 nM) on DY_m_ in ISO and A382T MNs 2 h after treatment at early and late time points. **(e)** Longitudinal study from a single MN culture comparing burst rate with DY_m_ measured at three time points. Data are shown as mean ± SEM. Statistical significance was determined using an unpaired t-test or a Mann-Whitney U test for normally and non-normally distributed datasets (Late: ISO vs A382T; Early Olig: ISO vs A382T; all within ISO (± Olig and Early/Late) comparisons). *p < 0.05, **p < 0.01, ***p < 0.001. Scale bars = 5 µm.

Similar to ETC electron flow (Figure 3a), ΔΨ_m_ is modulated by neuronal activity. Blockade of action potentials by TTX caused a reduction of ΔΨ_m_ in A382T MNs to levels observed in ISO MNs **(Figure 4d)**. Conversely, stimulation of neuronal firing by the potassium channel blocker tetraethylammonium (TEA) **(Supplementary Figure 3c)** led to mitochondrial hyperpolarization in both ISO and A382T MNs and occluded the difference in ΔΨ_m_ observed during the hyperexcitable phase **(Figure 4d)**. The prominent effect of TTX, together with the reduced impact of TEA on ΔΨ_m_, demonstrates a constant state of mitochondrial hyperpolarization in early-stage A382T MNs (DiC 34), which dissipates as these neurons age.

Co-increase in ETC activity and ΔΨ_m_ typically results in a higher proton motive force across the IMM and, thus, a greater potential for Fo/F1 ATPase activation. To assess engagement of the Fo/F1 ATPase in live MNs, we acutely blocked proton influx through the Fo channel with oligomycin and measured the ensuing increase in ΔΨ_m_.

Early on (DiC 34), oligomycin induced a robust increase in ΔΨ_m_ in both ISO and A382T MNs, indicating substantial usage of the proton gradient for ATP synthesis **(Figure 4d)**. Although the oligomycin-induced hyperpolarization was similar between genotypes, ΔΨ_m_ remained significantly higher in A382T MNs, consistent with elevated proton motive force and engagement of the Fo/F1

ATPase to meet the energetic demand during their hyperactive state. In older MNs (DiC 60), the oligomycin response was significantly blunted in ISO and undetectable inA382T mitochondria, implying a progressive loss of mitochondrial coupling and/or ATP synthase activity, which was more pronounced in A382T MNs.

To capture the transition between mitochondrial hyperpolarization and depolarization, we conducted an additional longitudinal study comparing MN firing with ΔΨ_m_ measured at three time points. We observed, again, ΔΨ_m_ reversal in A382T MNs (relative to ISO MNs), between early (DiC 34) and late (DiC 60) stages. At the peak of hyperexcitability (DiC 40), ΔΨ_m_ was not significantly different across groups, identifying a tipping point in mitochondrial function in A382T MNs.

### A382T MNs are ultra-sensitive to mitochondrial poisons

To determine whether changes in mitochondrial bioenergetics translated into altered ATP availability, we quantified total cellular ATP levels under normal growth conditions, permitting ATP synthesis via both glycolysis and oxidative phosphorylation (Glyc/Oxphos), or in presence of 2-deoxyglucose and galactose, forcing reliance on oxidative phosphorylation alone (Oxphos). During the hyperexcitable phase (DiC 27-33), steady-state ATP levels in A382T MNs and ISO MNs were indistinguishable under both Glyc/Oxphos **(Figure 5a, Supplementary Figure 4a)** and Oxphos **(Figure 5b, Supplementary Figure 4b)** conditions, indicating similar energy levels despite elevated ETC and Fo/F1 ATPase activity.

**Figure 5.**
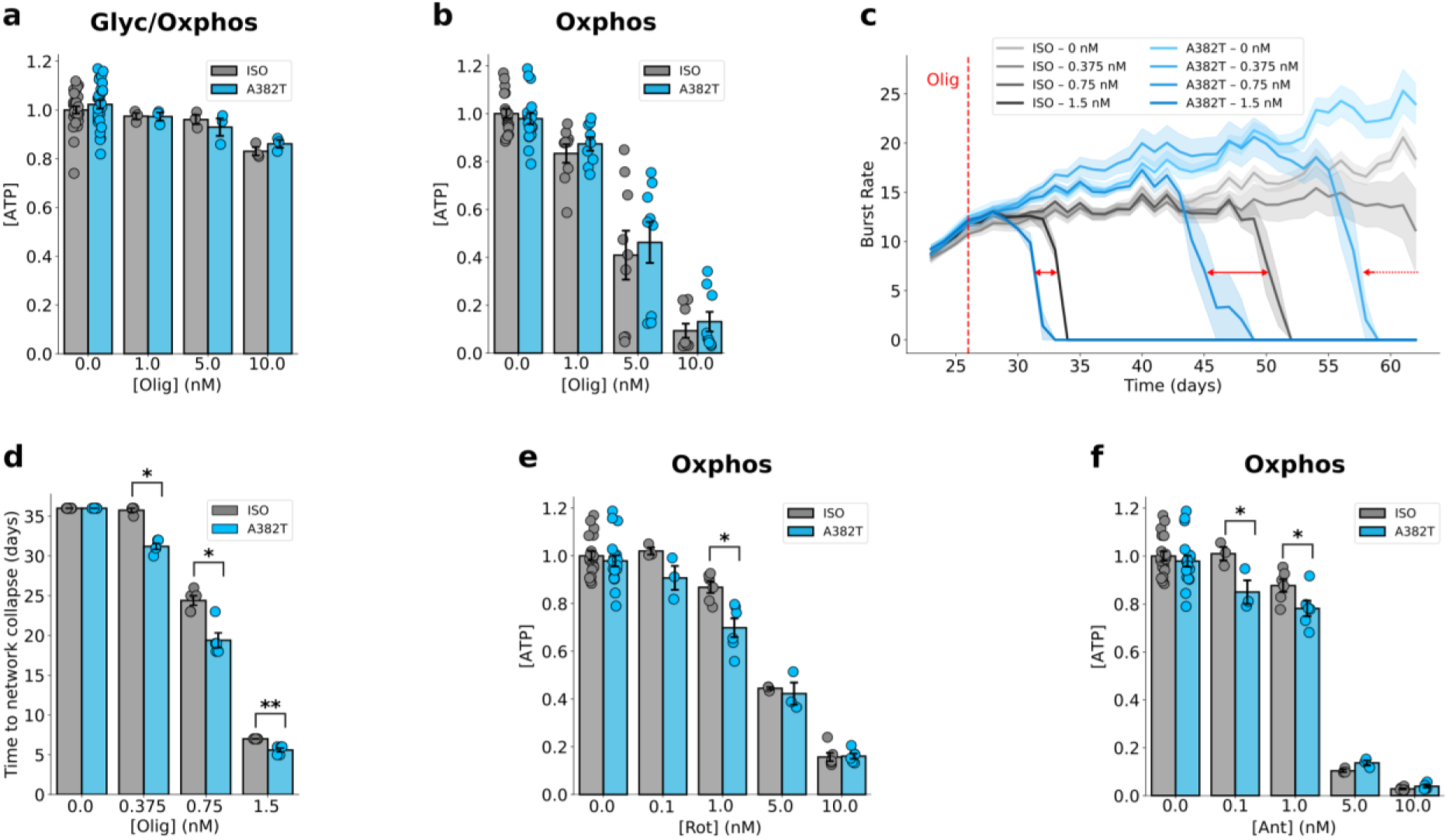
| A382T MNs are selectively vulnerable to mitochondrial stressors. (**a,b**) ATP measured using CellTiter-Glo under Glyc/Oxphos **(a)** or Oxphos **(b)** conditions, 4 h after exposure to vehicle or the indicated concentrations of oligomycin in ISO or A382T MNs at DiC 33. Data points represent replicate wells originating in **(a)** from 5 (vehicle) and 1 (remaining conditions) independent MN cultures and in **(b)** from 5 (vehicle) and 3 (remaining conditions) independent cultures. **(c)** Dose-dependent impact of oligomycin on burst rate in ISO (6 wells) and A382T (6 wells) MNs. The red dashed line indicates treatment onset, and red arrows compare ISO and A382T MNs for the same oligomycin concentration. **(d)** Comparison of the time to network collapse at different oligomycin concentrations. Data points represent individual wells (n > 4) originating from 1 independent MN culture. Statistical significance was tested using the Mann-Whitney U test, *p < 0.05, **p < 0.01. **(e,f)** Dose-dependent effect of rotenone **(e)** and antimycin A **(f)** on ATP levels 4 h after treatment of ISO and A382T MNs at DiC 33. Data points represent replicate wells from 2 (1 and 10 nM conditions) or 1 (remaining conditions) independent MN cultures. Data are shown as mean ± SEM. Statistical significance was assessed using the t-test or Mann-Whitney U test for normally and non-normally distributed datasets (Vehicle Glyc/Oxphos, Olig 5 & 10 nM Oxphos, and Rot 10 nM Oxphos) for genotype comparisons.

Short-term exposure (4 h) to increasing concentrations of oligomycin resulted in a dose-dependent decrease in ATP levels under Oxphos conditions, with near complete suppression of ATP at the highest dose in both ISO and A382T MNs **(Figure 5b)**. In contrast, ATP levels under Glyc/Oxphos conditions were largely maintained at all oligomycin concentrations **(Figure 5a)**, suggesting rapid and efficient metabolic rewiring towards glycolysis to compensate for loss of mitochondrial ATP. Early-stage ISO and A382T MNs were equally resistant in preserving cellular ATP levels at steady-state, in response to Fo/F1 ATPase inhibition **(Figure 5a,b)**. Together with the observed rise in ETC and Fo/F1 ATPase activity, these findings suggest that, early on, A382T MNs exhibit elevated ATP turnover to sustain higher firing rates, while maintaining energy homeostasis.

We reasoned that mitochondria in A382T MNs may operate near their bioenergetic limit to support increased ATP flux during the hyperexcitable phase. If so, partial inhibition of the Fo/F1 ATPase would exert additional pressure on these organelles and increase the risk of energy collapse with immediate consequences for neuronal firing. We therefore tested the dependence of neuronal firing to graded concentrations of oligomycin.

At a concentration ≥2.5 nM, oligomycin caused a rapid (<12 h) collapse of burst firing in both ISO and A382T MNs **(Supplementary Figure 4c)**, demonstrating a critical requirement for mitochondrial ATP synthesis. Although oligomycin significantly reduced mitochondrial ATP production **(Figure 5b)**, global ATP levels were largely preserved **(Figure 5a)**, suggesting that mitochondrial ATP is dispensable for steady-state energy maintenance, but becomes essential during periods of intense neuronal activity^4,8^.

Notably, lower concentrations of oligomycin (0.375-1.5 nM) differentially impacted firing in A382T and ISO MNs. These perturbations led to a dose– and time-dependent collapse of burst activity, with, for each concentration, a delay of several days for ISO MNs, indicating increased resilience to partial Fo/F1 ATPase inhibition **(Figure 5c, Supplementary Figure 4d,e)**. In A382T MNs, the increased sensitivity to mild Fo/F1 ATPase inhibition suggests that mitochondria function closer to their energetic ceiling, rendering them less able to compensate for modest deficits in ATP availability.

To further probe mitochondrial vulnerability to metabolic and oxidative stress, we exposed MNs to inhibitors of the ETC targeting complex I (rotenone) and complex III (antimycin A). These compounds limit (or suppress) electron flow and generate reactive oxygen species^57,58^. Both rotenone and antimycin A induce dose-dependent ATP depletion in Oxphos conditions, with A382T MNs displaying greater sensitivity to low concentrations (0.1-1 nM) than ISO controls **(Figure 5e, f)**, pointing to enhanced vulnerability of mitochondria to these ETC poisons. The toxicity of these inhibitors precluded long-term measurements of neuronal activity.

Together, these perturbation experiments demonstrate heightened vulnerability of A382T MNs to metabolic and oxidative stress, reflecting a state of pathological instability driven by near-maximal mitochondrial output during the hyperexcitable phase. In this regime, further increases in energetic demand cannot be accommodated, ultimately predisposing MNs to bioenergetic failure **(Figure 6)**.

**Figure 6.**
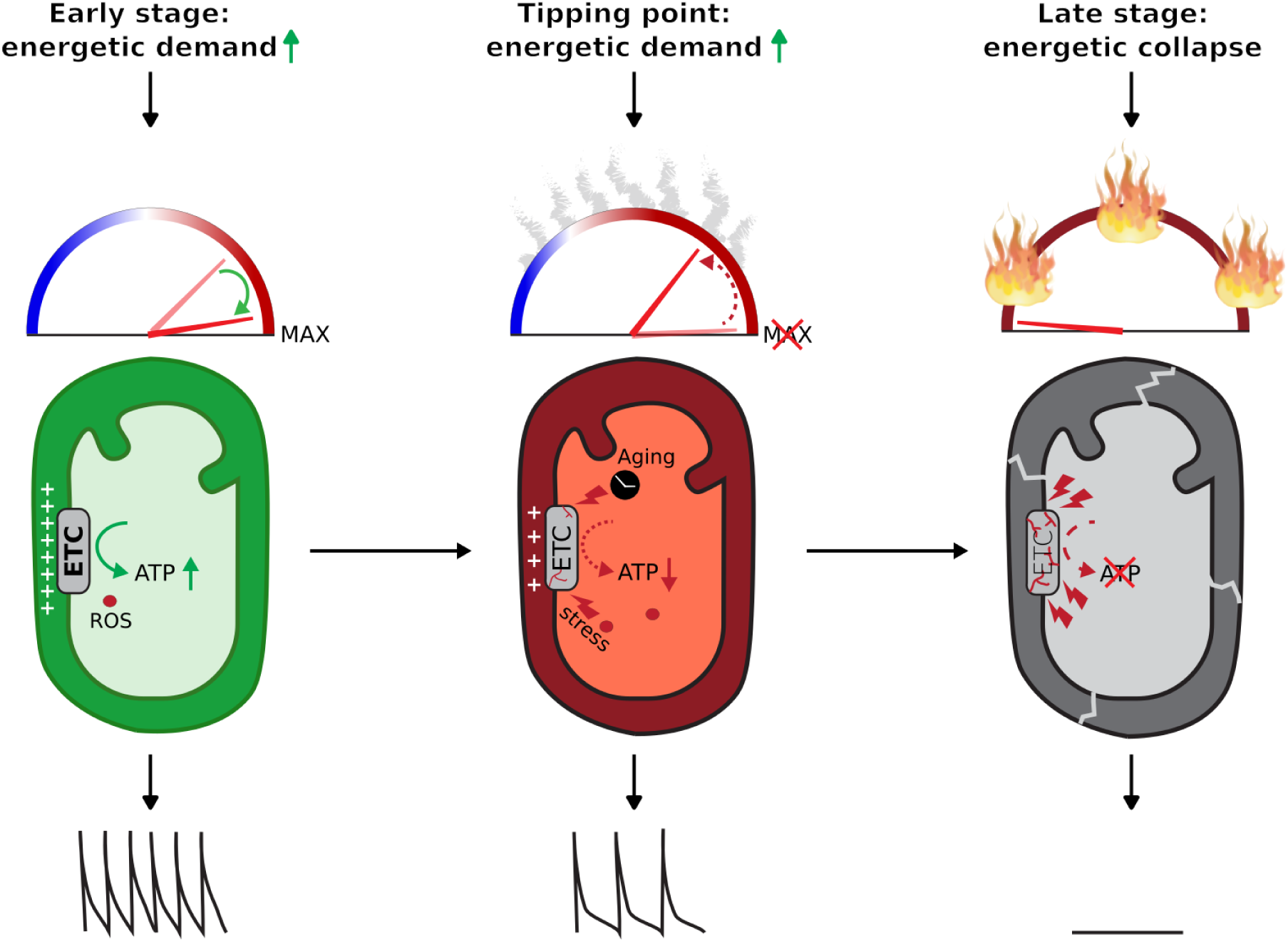
| Proposed model for ALS progression integrating pathological firing behavior with mitochondrial energy metabolism. Early on (DiC < 40), mitochondria operate near their bioenergetic limit to support increased neuronal activity. This prolonged near-maximal activity likely generates ROS, which gradually impair ETC and Fo/F1 ATPase functions. Accumulating oxidative stress pushes mitochondria beyond a critical tipping point (DiC 40-45), leading to ATP depletion, reduced neuronal output, and motor neuron degeneration. In late stages (DiC > 60), extensive mitochondrial damage results in bioenergetic collapse and loss of neuronal activity. Arrows indicate temporal progression from early hyperactive compensation to late-stage energy collapse.

## DISCUSSION

We describe high-resolution trajectories of spontaneous network activity in human iPSC-derived MN cultures expressing ALS TDP-43 variants, featuring an early, transient phase of hyperexcitability followed by progressive decline in firing parameters and loss of firing-competent MNs. This firing behavior is closely mirrored by changes in mitochondrial parameters, defining a tipping point in energy metabolism linking hyperexcitability to MN failure.

Acute manipulations of MN activity revealed tight coupling between MN firing rates and mitochondrial outputs and suggests that enhanced mitochondrial metabolism during the hyperexcitable phase is a compensatory response of A382T MNs to higher energy demands. In this transient state, both ETC activity and ΔΨ_m_ are elevated, consistent with enhanced respiration and tight coupling between oxidative phosphorylation and Fo/F1 ATPase output. Together, these data suggest prolonged engagement of A382T MNs in ADP-dependent (State III) respiration for enhanced production of ATP^59–61^.

Two additional lines of evidence support the notion that mitochondria in A382T MNs function near their energetic ceiling in the hyperexcitable phase. First, robust neuronal stimulation by TEA leads to a smaller ΔΨ_m_ increase, reflecting a hyperpolarized steady state, operating closer to that observed during intense, drug-induced, firing. Second, A382T MN firing is exquisitely sensitive to minor Fo/F1 ATPase inhibition. This heightened sensitivity indicates limited ability to compensate for minute ATP deficits and suggests that a large fraction of mitochondria operate near their bioenergetic limit.

This chronic, near-maximal mitochondrial engagement has long-term consequences for mitochondrial function and ALS progression. Elevated electron flow through respiratory complexes is typically associated with increased electron leakage and production of ROS^62,63^, which, in excess, can damage the ETC and the Fo/F1 ATPase^64,65^. Accordingly, low doses of the ETC poisons rotenone or antimycin A – known ROS producers – cause greater ATP depletion in A382T neurons, suggesting elevated susceptibility to oxidative stress or reduced redox buffering. Together, the limited resilience to both increased energetic demand and perturbation of the respiratory chain reveal two mechanistically linked stress sources that likely amplify age-dependent mitochondrial and neuronal decline in ALS^66–68^.

How is hyperexcitability coupled to mitochondrial bioenergetics? One prominent pathway involves activity-dependent calcium uptake, which boosts TCA cycle and ETC activity, thereby increasing ATP production^8,69^. However, sustained calcium influx can impair mitochondrial buffering capacity and trigger opening of the mitochondrial permeability transition pore, an important ALS therapeutic target^40,70^.

Based on these findings, we propose a disease model linking two key hallmarks of ALS: aberrant MN firing and disturbances in mitochondrial function. Early hyperexcitability drives chronic mitochondrial hypermetabolism, creating metabolic and oxidative stress that predispose A382T MNs to mitochondrial damage. As this stress accumulates with age^66–68^, mitochondrial resilience progressively declines until a critical “tipping point” is reached, precipitating bioenergetic failure and impaired firing **(Figure 6)**. Our findings further suggest that combined interventions aimed at enhancing mitochondrial resilience^71^, limiting oxidative stress^72^, and reducing excessive neuronal firing^73^ during the hyperactive phase may help preserve MN function and delay ALS progression.

## MATERIAL & METHODS

### iPSC Cultures

Induced pluripotent stem cell (iPSC) lines were obtained from Infinity Biologix (NINDS; NN0005319, homozygous reverted A382A; NN0005320, heterozygous A382T) and from The Jackson Laboratory (JIPSC001108, heterozygous M337V; JIPSC001110, homozygous reverted M337M). Cells were maintained on Matrigel^TM^ (Thermo Fisher Scientific, 11573560)-coated 6-well plates (Greiner, 657160). Matrigel^TM^ was thawed on ice, diluted in DMEM/F12 (Thermo Fisher Scientific, 31330-038) according to manufacturer’s instructions, and plates were incubated at 37 °C and 5% CO_2_ for 30 min before use. iPSCs were passaged at ∼80% confluence using ReLeSR^TM^ (STEMCELL Technologies, 100-0483) for up to four subsequent passages. Medium was changed daily using mTeSR^TM^1 (STEMCELL Technologies, 85850).

### Differentiation of Spinal MNs

iPSCs were grown to ∼80% confluence and dissociated with Accutase^TM^ (STEMCELL Technologies, 07920) to generate a single-cell suspension in DMEM/F12. Viable cells were counted using a NucleoCounter, and 10000-12500 cells were seeded in wells of a U-shaped 96-well plate (Greiner, 10638441) pretreated with Anti-Adherence Rinsing Solution (STEMCELL Technologies, 07010). Cells were cultured in Differentiation Medium 1: Basal Medium (DMEM-F12/ Neurobasal^TM^ (Thermo Fisher Scientific, 21103-049) 1:1 + 1x Pen/Strep (Thermo Fisher Scientific, 15070-063) + 1x N2 Supplement-A (STEMCELL Technologies, 07152) + 1x Neurocult^TM^ SM1 Without Vitamin A (STEMCELL Technologies, 05731) + 2 mM L-Glutamine (Thermo Fisher Scientific, 25030-24) + 0.5 µM L-Ascorbic Acid (Sigma-Aldrich, A5960) + 50 µM β-mercaptoethanol (Thermo Fisher Scientific, 11528926)) supplemented with 5 µM Y-27632 (Bio-Tocris, 1254) + 40 µM SB431542 (Tocris, 1614) + 0.2 µM LDN-193189 (Bio-Connect, 04-0074) + 3 µM CHIR99021 (Tocris, 4423). Plates were centrifuged for 3 min at 100 g to facilitate embryoid body formation and incubated at 37 °C and 5% CO_2_ (Day in Culture (DiC) –10).

Medium was partially replaced (2/3) with successive differentiation media as follows:

- DiC –9: Differentiation Medium 1
- DiC –8: Differentiation Medium 2.1: Basal Medium supplemented with 0.5 µM Retinoic Acid (R2625, Sigma-Aldrich) + 0.5 µM SAG (566660, Sigma-Aldrich) + 3 µM CHIR99021
- DiC –6: Differentiation Medium 2.2 (same as 2.1 but without CHIR99021)
- DiC –3: Differentiation Medium 3: Basal Medium supplemented with 0.5 µM Retinoic Acid, 0.5 µM SAG, 0.01 µg/mL hrBDNF (STEMCELL Technologies, 78005.1), and 0.01 µg/mL hrGDNF (STEMCELL Technologies, 78058.1)
- DiC –1: Differentiation Medium 4: Basal Medium supplemented with 0.5 µM Retinoic Acid, 0.5 µM SAG, 0.01 µg/mL hrBDNF, 0.01 µg/mL hrGDNF, and 10 µM DAPT (Tocris, 2634).

On DiC 0, spheres were dissociated in 0.05% Trypsin-EDTA (Thermo Fisher Scientific, 11580626) and replated at 50 000 – 100 000 cells/cm^2^ onto either a PhenoPlate (Revvity, 6055302) or a black µClear F-bottom 96-well plate (Greiner, 655096), coated with 15 µg/ml poly-L-ornithine (PLO; Sigma-Aldrich, P4957) and 10µg/ml Mouse Laminin (LAM; Sigma-Aldrich, L2020). Cells were allowed to settle for 10-15 min at room temperature and then placed at 37 °C and 5% CO_2_. Twenty-four hours later, ½ medium was replaced with Differentiation Medium 4.

Subsequent ½ medium changes were performed with:

- DiC 4: Differentiation Medium 5: Basal Medium supplemented with 0.01 µg/mL hrBDNF + 0.01 µg/mL hrGDNF + 20 µM DAPT.
- DiC 6: Differentiation Medium 6: Basal Medium supplemented with 0.01 µg/mL hrBDNF + 0.01 µg/mL hrGDNF + 20 µM DAPT + 0.01 µg/mL hrCNTF (STEMCELL Technologies, 78010).
- DiC 7, 8, and 10: Differentiation Medium 7: Basal Medium supplemented with 0.01 µg/mL hrBDNF + 0.01 µg/mL hrGDNF + 0.01 µg/mL hrCNTF.

From DiC 12 onwards, ½ medium changes occurred three times per week with BrainPhys Complete medium (BPC): BrainPhys^TM^ (STEMCELL Technologies, 05790) supplemented with 1X Pen/Strep, 1X N2 Supplement-A, 1x Neurocult^TM^ SM1 (STEMCELL Technologies, 05711), 200 nM L-ascorbic acid, and 1 µM dibutyryl cyclic AMP (Sigma-Aldrich, D0627) until the time of assay.

### Longitudinal Spontaneous Live-cell Calcium Transient Imaging

MNs seeded at a density of 75 000 or 100 000 cells/cm^2^ were transduced at DiC 20 with 2.5 µl Neuroburst^TM^ Orange lentivirus (Sartorius, 4736) per well to express a genetically encoded calcium sensor. Longitudinal live-cell calcium imaging was performed using the Incucyte® S3 live-cell imaging system (Sartorius) under standard cell culture conditions (37 °C, 5% CO_2_) for up to 50 days. At DiC 25 or DiC 35, cultures were treated three times a week (Mon, Wed, and Fri) with different concentrations of oligomycin A (Tocris, 4110) or vehicle (Dimethylsulfoxide; Sigma-Aldrich, 67-68-5) to assess the effects of chronic mitochondrial ATPase inhibition. Spontaneous calcium transients were recorded daily for 1 minute per well at a frame rate of 3 frames per second. Calcium activity parameters, including active object count (active objects), burst rate, burst duration, burst strength, and the correlation (network correlation), were quantified using the built-in Neuronal Activity Analysis Software Module (Sartorius). For chronic treatment studies, transduced MNs were treated with oligomycin A starting between DiC 27-35 and were subsequently re-treated every Monday, Tuesday, and Friday until the end of the experiment. Data were further analyzed in python to generate t-SNE maps of the extracted parameters and to calculate the day after treatment where the active object count reached a value of 100 objects/well, once this point was reached, the burst rates following this value were also transformed to zero, to account for residual noise from the surviving active objects. If no threshold was reached, the maximum experimental time point was used. To accurately calculate the network collapse day after treatment, burst activity values were smoothed with a rolling window of 3 (pandas.Series.rolling) to minimize noise.

### High-density Microelectrode Array (MEA) Recordings

At DiC 0 of differentiation, progenitor cells from both lines were dissociated using Accutase^TM^ and plated onto a sterile CorePlate™ (MEA; 3Brain, 24W 16/50) pre-coated with 100 µg/mL PLO and 50 µg/mL LAM at a density of 80 000 cells per MEA. Differentiation was continued directly on the MEA plates under standard culture conditions (37 °C, 5% CO_2_). At DiC 25, recordings were performed using a BioCAM Duplex system (3Brain) at 37 °C for 3 min per MEA using Brainwave software (3Brain). Raw electrophysiological data were exported to Python for further analysis. Custom Python scripts were used to quantify neuronal network activity parameters. A spike was defined as the activation of a single electrode, a burst as >3 consecutive spikes on a single electrode with interspike intervals <0.1 s, and a burst wave as a network event in which >10 electrodes exhibit bursting activity within a 0.1 sec window. From these events, the following parameters were extracted: the spike rate (spikes/min), inter-spike interval (ISI; sec), the number of active electrodes (Active ch.), burst duration (Burst dur.; sec), bursts per channel per minute (Burst rate), the spikes per burst (Spk/burst), the burst magnitude (Burst mag.; Burst rate x Burst dur.), burst wave duration (Wave dur.; sec), the number of channels active per wave (Ch/wave), and the wave magnitude (Wave mag.; Wave dur. x Ch/wave). All extracted parameters were summarized as log_2_ fold changes (A382T/ISO), and spikes per burst were further visualized in a cumulative probability plot to highlight more subtle differences. Both genotypes were represented by three MEAs, all acquired from the same differentiation batch.

### Electron Flow Assay

Custom S-1 MitoPlates^TM^ (Biolog) coated with either purified water (no substrate), L-malic acid (Mal), succinic acid (Succ), α-ketoglutaric acid (αKG), tryptamine, pyruvic acid supplemented with a 100 µM spike of Malic Acid (Pyr + trace Mal), Glutamine, or Isocitric Acid (substrate concentrations proprietary to Biolog) were designed upon request. Substrates that did not yield a measurable signal above background (no substrate, tryptamine, glutamine, and isocitric acid) were excluded from analysis. Each substrate was pre-incubated for 30 min at 37 °C in the Biolog Mitochondrial Assay Mix: 1x Biolog Mitochondrial Assay Solution (Biolog, 72303) + 1x Redox Dye MC (Biolog, 74353) + 30 µg/mL saponin (Sigma-Aldrich, SAE0073) + distilled water to ensure complete dissolution. Following incubation, the culture medium was removed from the motor neuron plate, and 120 µL of the respective substrate solution was added to the assay plate (PhenoPlate) containing the neurons. Prior to kinetic measurements, a 4x image of each well was acquired using the Incucyte® S3 imaging system to assess neuronal coverage. Plates were transferred to a pre-heated (37 °C) Tecan M200 Pro plate reader, and absorbance was measured every 10 min at 590 and 720 nm (gain = 850, bandwidth = 90%, read height = 2 mm) for 3 hours. Experiments were conducted on early-stage (DiC 34) and late-stage (DiC 60) motor neurons, pre-treated for 24 h with either BPC (vehicle) or 1 mM tetrodotoxin citrate (TTX; Sanbio, SIH-603). Raw absorbance data were processed in Python. To correct for abiotic background, 720 nm absorbance values were subtracted from corresponding 590 nm readings for each well. Incucyte images were analyzed to quantify neuronal pixel coverage per well. Corrected absorbance values were then normalized to the pixel coverage x 1 000 000. The resulting normalized traces were then used to calculate the slope of the signal increase over the entire timeseries as a proxy for ETC activity. Due to differences in permeabilization efficiency, all individual substrates in each experiment were normalized to the slope of their respective ISO vehicle condition. This prevented comparisons across plates with different DiC.

### Mitochondrial Membrane Potential and Mitochondrial Abundance

Motor neurons were plated at densities ranging from 50 000 to 100 000 cells/cm2 in 96-well black PhenoPlates and assayed for mitochondrial membrane potential (ΔΨ_m_) at early (DiC 34), peak f hyperexcitability (DiC 40), or late (DiC 60) stages of maturation. Cells were treated with BPC (vehicle) or 4 mM tetraethylammonium chloride (TEA; Tocris, 3068), and four hours prior to imaging, additionally with 1 mM TTX or vehicle. Treatments were prepared in BPC supplemented with 20 nM tetramethylrhodamine methyl ester (TMRM; Thermo Fisher Scientific, T668). After two hours of TMRM incubation at 37 °C and 5% CO_2_, live-cell confocal imaging was performed on a PerkinElmer Operetta CLS using a 40x water objective (binning 2, excitation 530-560 nm, emission 570-650 nm, exposure 60 ms, laser power 40%). Six fields of views (FOVs) were acquired per well using a z-stack of 8 planes covering a depth of 14 µm. Image analysis was conducted in Harmony 5.2 (PerkinElmer). Background subtraction was performed using a sliding parabola (curvature = 1), after which individual mitochondria were segmented based on TMRM fluorescence using the Find Spots building block. Objects between 7 and 350 px^2^ that did not cross the edge of the FOV were included. Cell somata were detected by smoothing the background-subtracted TMRM signal and applying the Find Cells module. Morphological features (area, roundness, and width/length ratio) and TMRM intensity were used to train a machine-learning classifier to identify soma clusters. These clusters were then excluded to restrict analysis to mitochondria located in the neurite region. For each well, the mean TMRM intensity of all segmented mitochondrial objects was computed two hours after dye loading, serving as a proxy for mitochondrial membrane potential. Additionally, the total number of segmented TMRM-positive objects per well was used as a measure of mitochondrial abundance. Single-well data were exported to Python, where wells displaying morphological aberrations or z-score standard deviation >3 were excluded. Experiments were pooled based on their respective DiC.

### ATP Quantification

To assess steady-state ATP levels in MNs, cells were plated at a density of 100 000 cells/cm² on PLO/LAM-coated white F-bottom microplates (Greiner, 655074). At DiC 30, MNs received a ½ medium change with either BPC (vehicle) or BPC supplemented with 2.5 mM 2-deoxy-D-glucose (2-DG; Sigma-Aldrich, D8375) and 2.5 mM galactose (Sigma-Aldrich, G5388) to inhibit glycolysis and force MNs to rely on OXPHOS for ATP production. Both conditions were combined with a range of mitochondrial inhibitors – oligomycin A (olig), antimycin A (Ant; Sigma-Aldrich, A8674), rotenone (Rot; Tocris, 3616/50) – or DMSO (vehicle) to modulate mitochondrial functioning. After 4 h of treatment, the respective medium was removed, and cells were washed with PBS. Subsequently, CellTiter-Glo® detection reagent (Promega, G7570) was added directly to the neurons, together with an ATP standard curve (Sigma-Aldrich, A1852), and the plate was shaken for 2 min to induce cell lysis. After a 10 min incubation at room temperature in the dark, luminescence was measured using a Tecan M200 Pro microplate reader with an integration time of 0.25-1 s per well. To normalize ATP levels to cell number, an equal volume of lactate dehydrogenase (LDH) reagent from the CytoTox 96® Cytotoxicity Assay (Promega, G1780) was added to each well immediately after luminescence readout. LDH absorbance was measured at 492 nm every 5 min for 30 min using the same plate reader. ATP concentrations were calculated by fitting luminescence readings to the linear regression of the ATP standard curve. The resulting ATP concentration values were then divided by the background-subtracted LDH absorbance to yield [ATP]/cell. Data were exported to Python for further analysis. For each experiment, [ATP]/cell values were normalized to the mean ISO vehicle values under both standard ([ATP] from Glyc/Oxphos) and Oxphos-driven conditions ([ATP] from Oxphos), yielding the final metrics for inter-experimental comparisons.

### Statistical Analysis

All data are presented as mean ± SEM. Statistical analyses were performed using Python (v3.10; SciPy.stats). For TMRM measurements, Glyc/Oxphos and Oxphos ATP quantification, and electron flow assays, the individual well was used as the unit of analysis. The number of independent experiments per condition varied between 1 and 6 depending on treatment and concentration (see figure legend). Normality of distributions was assessed using the Shapiro-Wilk test. For normally distributed data, comparisons between two groups were performed using unpaired or paired t-tests, as appropriate; for non-normally distributed data, the Mann–Whitney U test or Wilcoxon signed-rank test was used. Multiple comparisons were corrected using the Benjamini-Hochberg false discovery rate procedure, when applicable. For calcium imaging experiments using Neuroburst^TM^, data represent individual independent experiments. Normality was assessed as above. Comparisons between two groups were performed using unpaired t-tests for normally distributed data and Mann–Whitney U tests for non-normally distributed data. Longitudinal analyses of neuronal activity were performed using the same statistical approach. No data points were excluded unless explicitly stated (e.g., failed wells or technical artifacts). All statistical tests are two-tailed unless otherwise indicated. A p-value < 0.05 was considered statistically significant. Exact p-values are reported in figure legends.

## Data availability

The datasets generated and analyzed during the current study, including raw and processed imaging data, analysis scripts, and final figures, are available in the Zenodo repository (https://doi.org/10.5281/zenodo.19221000). Data were generated at ReMYND and may be subject to access and reuse restrictions.

## Acknowledgements

This work was supported by Flanders Innovation & Entrepreneurship (VLAIO), formerly known as the Institute for the Promotion of Innovation by Science and Technology in Flanders (grants HBC.2021.0610; HBC.2022.0958). We thank ReMYND NV for its support. We are grateful to Kristel Eggermont of the Lab of Neurobiology (VIB, Leuven) for providing the differentiation protocol. We also thank Bary Bochner and Enrico Tatti for their contributions to the design of the custom Biolog plates and for their guidance in performing these experiments.

## Author contributions

J.P. designed and performed experiments, analyzed the data, contributed to the conceptual framework of the study, and wrote the original draft of the manuscript. T.V., M.v.G., S.V., and K.Pi. performed experiments and contributed to data acquisition and methodology development. T.L., I.B., and M.R. assisted with experimental work and data acquisition. K.Pr. contributed to conceptual input. M.v.G. contributed to figure preparation and manuscript editing. M.F. conceived and supervised the study and contributed to manuscript writing and editing. G.G. conceived and supervised the study and contributed to manuscript editing. F.S. contributed to supervision and funding acquisition. All authors reviewed and approved the final manuscript.

## Declaration of generative AI and AI-assisted technologies in the writing process

During the preparation of this manuscript, the authors used ChatGPT (OpenAI) to improve the clarity and readability of the text. After using this tool, the authors reviewed and edited the content as needed and take full responsibility for the content of the published article.

