## Supplementary Figures for "A mitochondrial tipping point couples early hyperexcitability to late-stage failure in patient-derived ALS motor neurons"

[1]reMYND NV; Bio-Incubator, Gaston Geenslaan 1, Leuven-Heverlee 3001, Belgium.

[2]Lead Contact

### SUPPLEMENTARY FIGURES

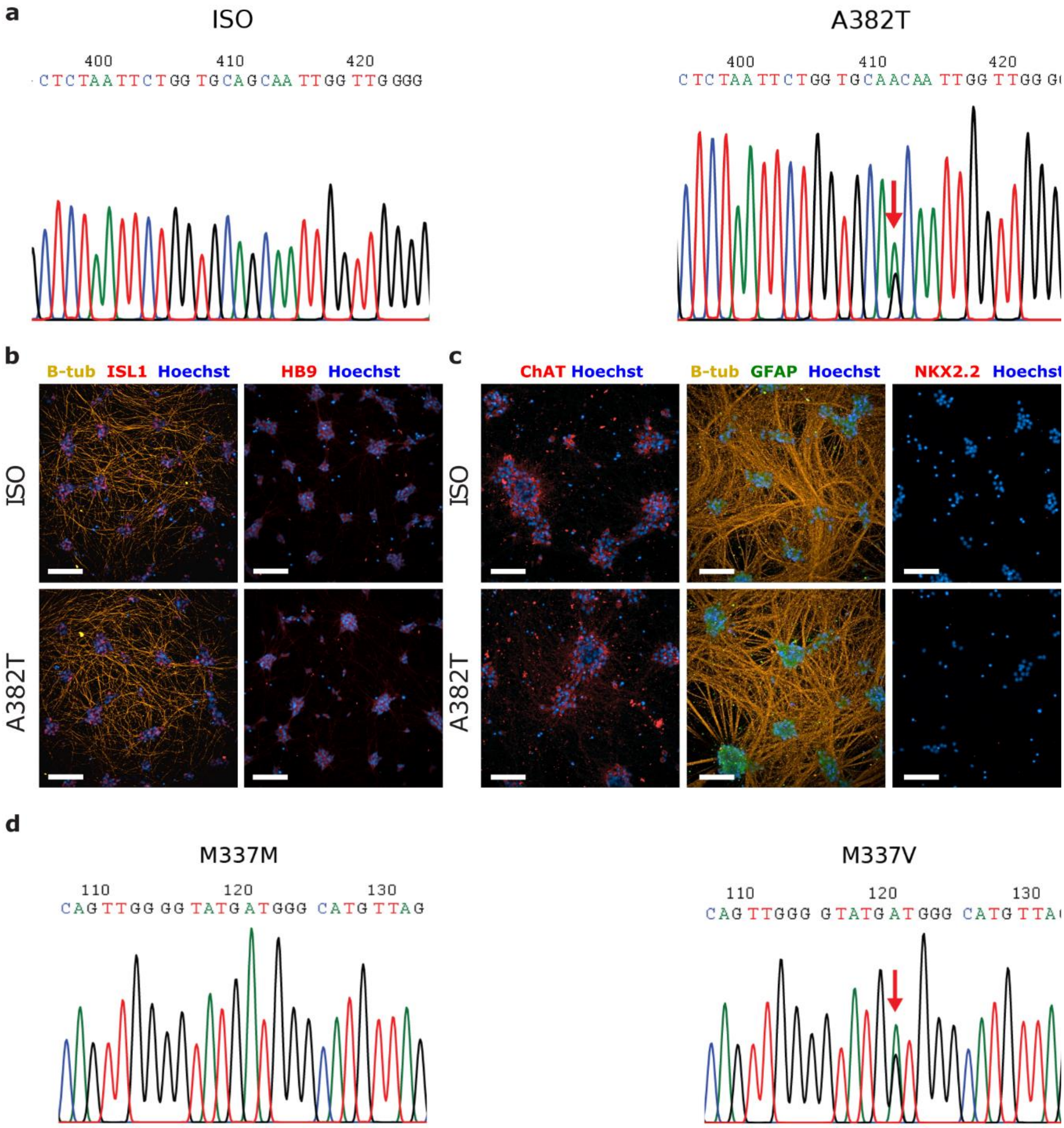


**Supplementary Figure 1 | Characterization of iPSC-derived MNs carrying the TDP-43 A382T mutation. (a)** Sanger sequencing confirming the presence of the heterozygous A382T mutation in TARDBP in patient-derived iPSCs and its reverted control (ISO). **(b)** Immunofluorescence characterization of motor neuron progenitors at DiC 4 showing expression of ISL1 (Islet-1) and HB9 (MNX1). **(c)** Mature MNs (DiC 40) stained for choline acetyltransferase (ChAT), glial fibrillary acidic protein (GFAP) and NKX2.2 (interneuron marker) protein. **(d)** Sanger sequencing confirming the presence of the heterozygous M337V mutation in TARDBP in patient-derived iPSCs and its reverted control (ISO).

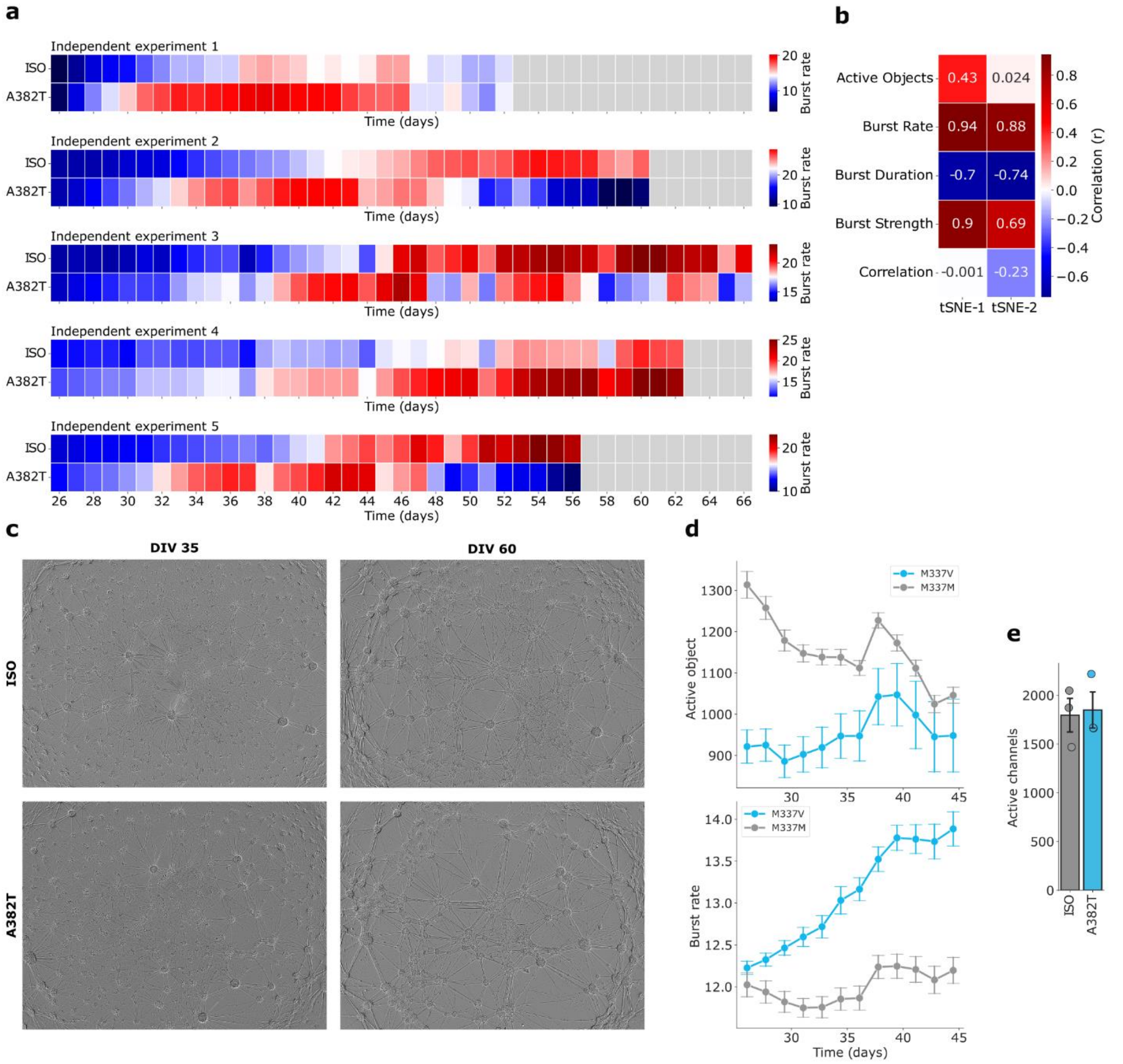


**Supplementary Figure 2 | Neuronal activity readouts in A382T and M337V MNs. (a)** Heatmaps of all experiments that reached DiC 50, illustrating the longitudinal change in burst rate for ISO (n = 6 wells) and A382T (n = 6 wells) neurons. **(b)** Feature–tSNE correlation heatmap illustrating how individual firing parameters relate to the tSNE-1 and tSNE-2 dimensions. Positive and negative correlations indicate the relative contribution of each parameter to the spatial distribution of data points. **(c)** Representative Incucyte image (4x) of early (DiC 35) and late-stage (DiC 60) motor neurons for both ISO and A382T genotypes. **(d)** Multidimensional analysis of spontaneous neuronal firing in M337V and ISO TDP-43 MNs from DiC 25 to DiC 45. Time series showing active object count and burst rate (mean ± SEM) from averaged wells originating from 1 independent MN culture. **(e)** Quantification of the number of active MEA electrodes (Active channels) during the readout period at DiC 25. Values represent averaged electrodes from 3 independent MEAs originating from the same differentiation.

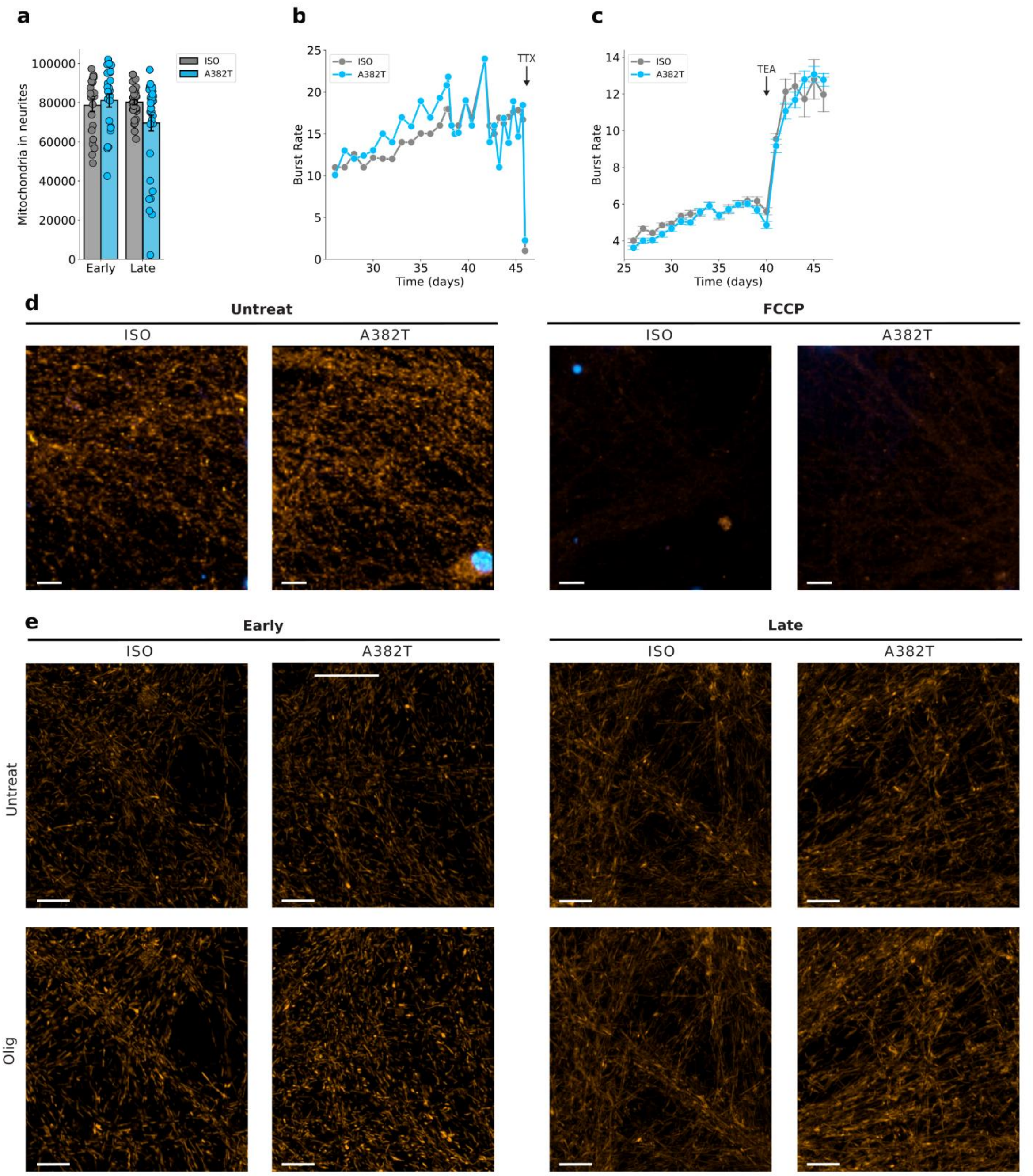


**Supplementary Figure 3 | Effect of neuronal and metabolic modulators on neuronal activity and mitochondrial membrane potential. (a)** Amount of TMRM-positive mitochondria in neurites as an indication of viable mitochondria at early (DiC 34) and late (DiC 60) stages in ISO and A382T MNs. **(b,c)** Effect of neuronal modulators on burst rate, with TTX silencing and TEA increasing neuronal activity. TTX data were obtained from individual wells, and TEA data were obtained from 3 averaged wells originating from 1 independent experiment with a low burst rate. Values represent mean ± SEM. **(d)** $\Delta\Psi_m$ imaged in MNs treated for 1 h with vehicle or 5 μM FCCP. **(e)** $\Delta\Psi_m$ imaged by TMRM staining in neurites of ISO and A382T MNs at early (DiC 34) and late (DiC 60) time points right before treatment (untreat) and 2h after treatment with oligomycin (Olig). Scale bars = 5 μm.

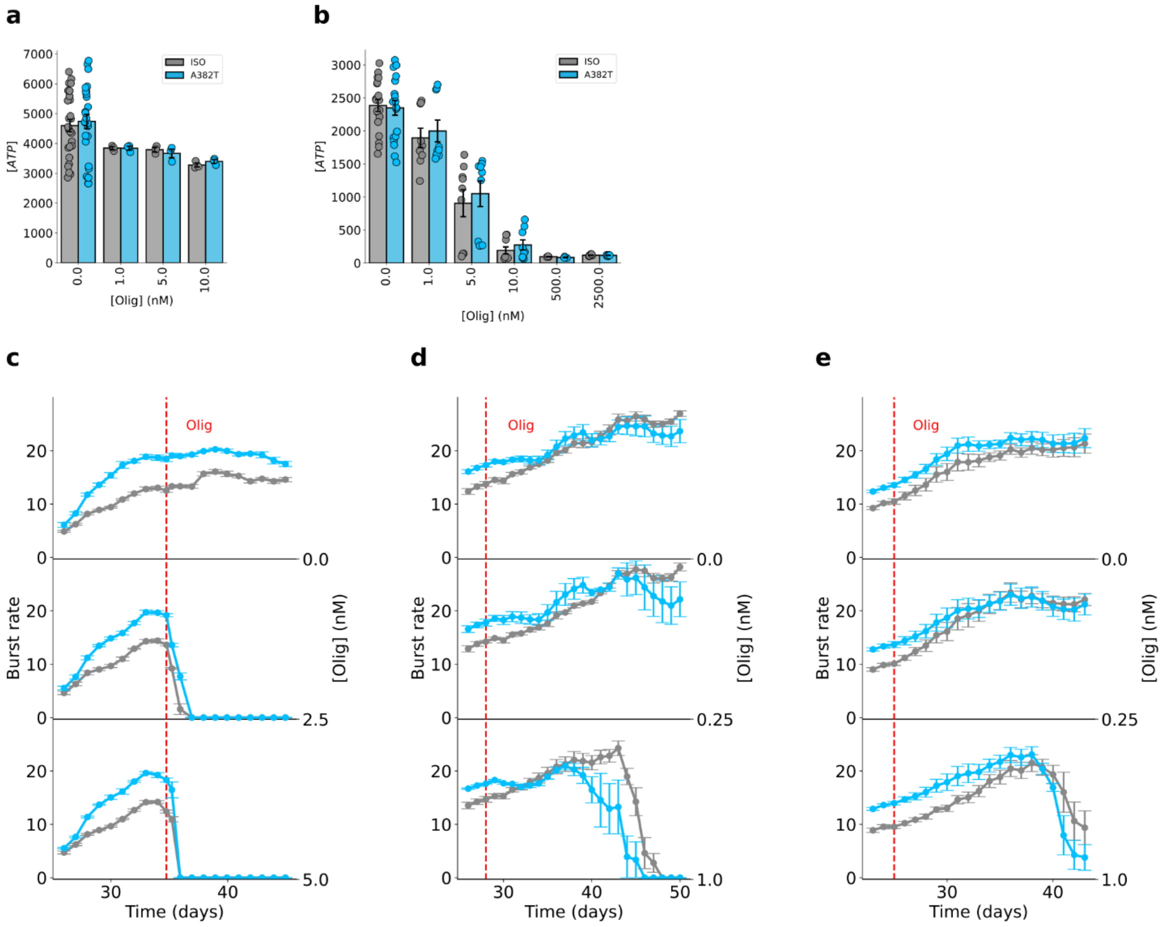


**Supplementary Figure 4 | Sensitivity of motor neuron activity to mitochondrial stressors is reproduced in independent experiments. (a,b)** Raw CellTiter-Glo values, normalized to their respective LDH values per well, for both ISO and A382T cells in Glyc/Oxphos **(a)** and Oxphos-only conditions **(b)**. **(c)** Representative ridge plot of neuronal burst rate over time during late early-stage (DiC 35) chronic exposure to high oligomycin concentrations. The red dashed line indicates the onset of oligomycin treatment. (**d,e**) Representative ridge plots of 2 independent experiments chronically treated with low concentrations of oligomycin starting from an early time point (DiC < 30). All data points represent individual wells from their respective independent experiments. Values represent mean ± SEM.
